# A synthetic Notch (synNotch) system linking intratumoral immune-cancer cell communication to a synthetic blood biomarker assay

**DOI:** 10.1101/2023.09.26.559329

**Authors:** YangHao Fu, TianDuo Wang, John A. Ronald

## Abstract

**Introduction:** Cellular immunotherapy has greatly improved cancer treatment in recent years. For instance, chimeric antigen receptor (CAR) T-cell therapy has been proven highly effective in treating hematological malignancies, and many CAR cell designs are being explored for solid tumors. However, many questions remain why responses differ across patients and some tumor types are resistant. Improved and relatively inexpensive ways to monitor these cells could provide some answers. Clinically, blood tests are regularly used to monitor these therapies, but blood signals often do not reflect the activity of immune cells within the tumor(s). Here, using the synthetic Notch (synNotch) receptor that tethers antigen binding to customized transgene expression, we linked intratumoral immune-cancer cell communication to a simple secreted reporter blood test. Specifically, we engineered immune cells with a CD19-targeted synNotch receptor and demonstrate that binding to CD19 on cancer cells in vivo resulted in the production of secreted embryonic alkaline phosphatase (SEAP) at levels that are readily detected in the blood.

**Methods and Results:** Jurkat T cells were engineered via sequential lentiviral transduction of two components: an anti-CD19 synNotch receptor and asynNotchresponse element encoding SEAP. Co-culture of engineered cells with CD19^+^, but not CD19^-^, Nalm6 cells, resulted in significantly elevated SEAP in media. Nod-scid-gamma (NSG) mice were subcutaneously injected with either CD19^+^ or CD19^-^ Nalm6 cells. Intratumoral injection of engineered T cells (1x10^7^) resulted significantly elevated blood SEAP activity in mice bearing CD19^+^ tumors (n=7), but not CD19^-^ tumors (n=5).

**Discussion:** Our synNotch reporter system allows for the monitoring of antigen-dependent intratumoral immune-cancer cell interactions through a simple and convenient blood test. Continued development of this system for different target antigens of interest should provide a broadly applicable platform for improved monitoring of many cell-based immunotherapies during their initial development and clinical translation, ultimately improving our understanding of design considerations and patient-specific responses.

## INTRODUCTION

Cell-cell communication plays a vital role in human development, homeostasis, and pathogenesis^1–3^. The advent of therapeutic cells specifically designed to interact and communicate with diseased cells has ushered in a new era in the treatment of numerous medical conditions. This approach has already shown promising results for infectious diseases^4^, immunologic deficiency syndromes^5,6^, neurodegenerative and movement disorders^7^, as well as cancer^8,9^.

T cells have been extensively used in cell-based cancer immunotherapies due to their cytotoxic capabilities, and the ability to home to and proliferate within tumors upon adoptive transfer^10–13^. Specifically, T cells can elicit cytotoxic effects through interactions between endogenous or engineered receptors with molecular targets on cancer cells^14,15^. Although these therapies have been transformative for the treatment of numerous malignancies, their ineffectiveness in some individuals often stem from poor tumor homing, a hostile tumor microenvironment, tumor heterogeneity, and antigen loss or escape^16,17^. Moreover, on-target/off-tumor toxicities can sometimes result in detrimental and even life-threatening side effects^18–22^. To better understand the effects of these therapies in individual patients, assays to monitor therapeutic cells over time would be of great benefit to develop safer and more robust immunotherapies.

Affordable and minimally invasive blood assays offer convenient ways to monitor *in vivo* biological events. For instance, blood tests using quantitative PCR or flow cytometry have been used to directly monitor the levels and persistence of adoptively transferred immune cells over time^23^. However, there is often a disconnect between circulating blood measures of immune cell numbers and how many there are in tumors, the site of action. For instance, quantitative measures of intratumoral chimeric antigen receptor T (CAR-T) cells using PET reporter gene imaging was shown to not correlate with blood levels of CAR-T cells in mice^24^. Moreover, current blood assays do not easily reflect the activity and behavior of intratumoral T cells. Here we sought to develop an assay that links antigen-specific T cell interactions with cancer cells within tumors to an easily measurable synthetic biomarker in the blood. Specifically, we leveraged an activatable synthetic biology system called the synthetic Notch (synNotch) receptor to relay intratumoral immune cell-cancer cell communications into the blood activity levels of a secreted reporter gene.

The synNotch system was described in a series of papers in 2016 as a general platform for building novel cell-cell contact signaling pathways^25–27^. This system is composed of both the synNotch receptor and the synNotch response element (RE), which are co-engineered into the same cell. Traditionally, like clinically-used CARs, the extracellular domain of the synNotch receptor is comprised of the single-chain variable fragment (scFv). Upon binding to the target antigen, this interaction triggers the cleavage of the GAL4-VP64 intracellular domain of the receptor. Subsequently, the GAL4-VP64 fusion protein binds to the upstream activation sequences (UAS) located within a minimal promoter. This binding event leads to the activation of transcription, thereby driving the expression of transgenes of interest encoded in the RE. By maintaining the Notch core regulatory region but appending a customized extracellular input recognition and intracellular output transcription activator module, novel cell-to-cell contact signaling pathways that carry out user-defined functionalities can be established^25^. We have recently linked synNotch antigen binding with the transcriptional activation of imaging reporter genes to monitor immune cell-cancer cell communication *in vivo* with both bioluminescence and clinically relevant magnetic resonance imaging^28^. Similarly, a next-gen synNotch system has been linked to positron emission tomography (PET) reporter gene expression and antigen-activation has been detected using PET^29^. However, while imaging provides important spatial information, the ability to perform repetitive imaging can be expensive, especially in patients. To complement our imaging system, we developed a synNotch system where the output is a convenient blood assay using a human-derived secreted reporter gene called secreted embryonic alkaline phosphatase (SEAP) (Figure 1A).

**Figure 1.**
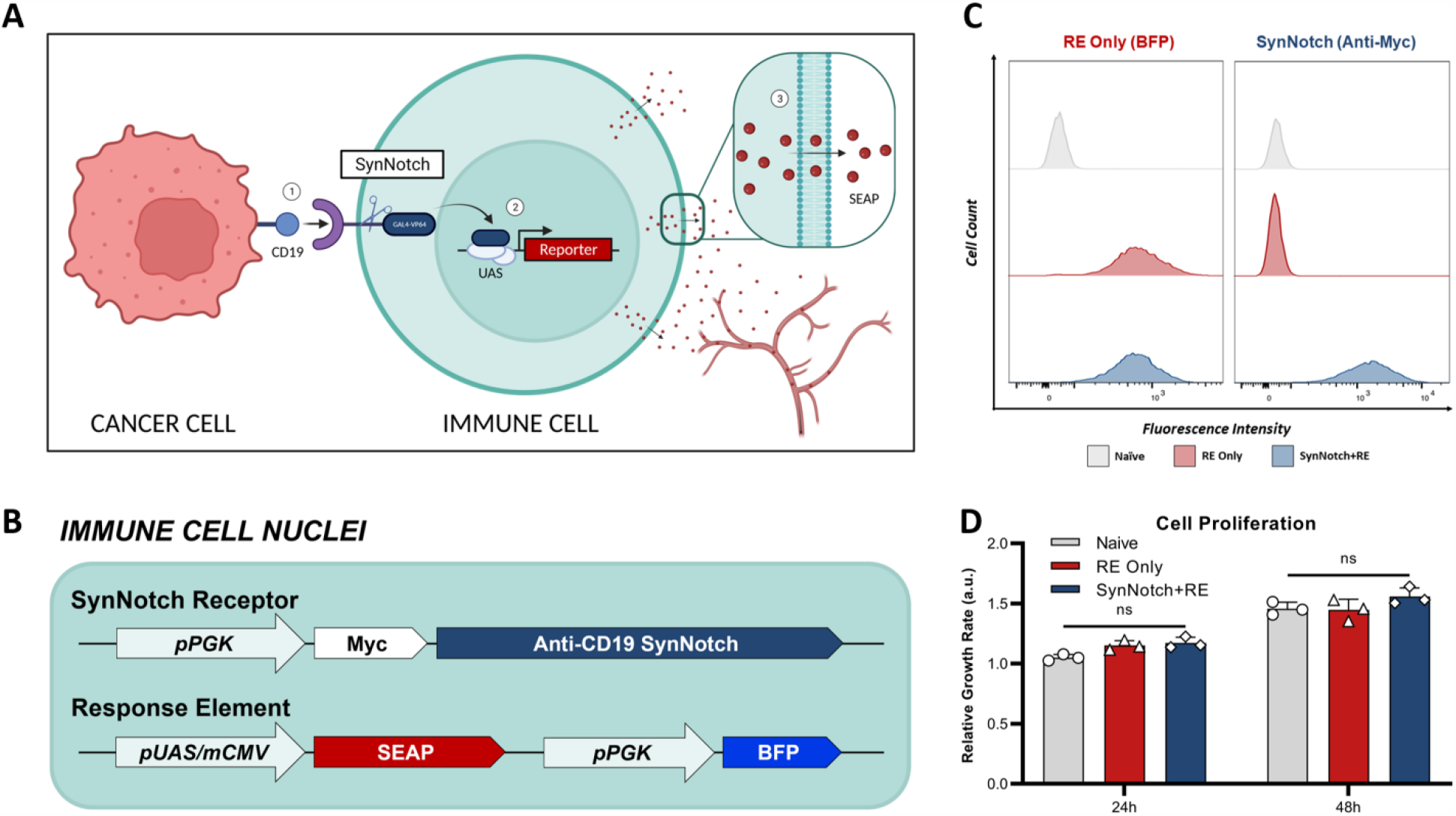
Engineering of T cells with blood-based reporter system for cell-cell communication. **(A)** Schematic of synNotch system. The synNotch receptor has three components: the an antigen-targeting extracellular domain; a core regulatory region that controls cleavage of the receptor upon antigen interaction; and an intracellular domain comprised of a GAL4-VP64 DNA-binding/transcriptional activator fusion protein^26^. Upon binding of the anti-CD19 synNotch receptor in T cells to CD19 on an adjacent cancer cell, GAL4-VP64 is cleaved from the synNotch receptor and can bind the upstream activation sequence (UAS) of a minimal promoter in the response element (RE) to drive expression of transgenes of interest^52^ – such as secreted embryonic alkaline phosphatase (SEAP). *In vivo*, SEAP can diffuse into the bloodstream and measured using a luminescence-based assay in blood samples. **(B)** T cells were first transduced with a response element (RE) containing SEAP and a constitutively expressed blue fluorescence protein (BFP) driven by the phosphoglycerate kinase 1 promoter (pPGK) for cell sorting. (**C**) Flow cytometry of naïve and twice transduced T cells post-sorting, assessing the expression of RE (BFP^+^) and synNotch (Myc+). **(D)** Relative growth rates of T cell populations at 24- and 48-hours post-seeding (n=3). Data presented as mean ± SD.

SEAP stands out as the most widely utilized secretable reporter protein due to its distinct characteristics. It is a truncated form of human placental alkaline phosphatase that is expressed specifically during embryogenesis, with minimal post-natal expression^30–32^, rendering it a highly specific blood reporter. SEAP exhibits heat stability, specifically, the heating of serum samples to 65°C enables its selective detection in assays without detecting other phosphatases that may be present in the blood^33,34^. Commercial SEAP detection assays are also extremely sensitive over at least a 4-log order concentration range, with detection limits in the picogram per milliliter range. Additionally, murine SEAP showed no immunogenic potential in mice, implying that human-derived SEAP is safe for clinical translation, and has already been successfully used in the clinic^34,35^. For the antigen target of our synNotch receptor we chose CD19 as a proof-of-concept due to its effectiveness as a target for CAR-T cell therapies against B cell leukemia and lymphomas^36,37^.

In this study we developed and validated a novel synNotch system to provide specific readouts of intratumoral CD19-triggered immune-cancer cell communication by measuring blood SEAP activity levels. Continued development of this cell-cell communication blood assay, and merger with other readouts such as imaging, has the potential to provide extended insight on the efficacy of T cell immunotherapy during both its development and clinical integration.

## MATERIALS AND METHODS

### Lentiviral Design and Production

A lentiviral transfer plasmid encoding encoding an anti-CD19 synNotch receptor driven by pPGK was acquired from Addgene (pHR_PGK_antiCD19_synNotch_Gal4VP64 was a gift from Wendell Lim; Addgene plasmid # 79125; http://n2t.net/addgene:79125; RRID:Addgene_79125).

A response element (RE) construct was made by also acquiring a lentiviral transfer plasmid containing the GAL4-VP64 inducible upstream activator sequence (UAS) from Addgene (pHR_5x Gal4 UAS was a gift from Wendell Lim (Addgene plasmid # 79119; http://n2t.net/addgene:79119; RRID:Addgene_79119). The SEAP transgene from our previously developed plasmid pSurvivin-SEAP-WPRE (REF) was cloned into pHR_5x GAL4 UAS using In-Fusion HD cloning (Takara Bio, CA, USA) to make pHR_5x Gal4 UAS SEAP. A constitutively expressed blue fluorescence protein (BFP) driven by the phosphoglycerate kinase promoter (pPGK) was inserted further downstream of the same reporter cassette via the same cloning kit to make pHR_5x Gal4 UAS SEAP pPGK BFP.

A second-generation lentiviral packaging plasmid (pCMV delta R8.2, #12263), and envelope plasmid (pMD2.G, #12259) were acquired from Addgene (pCMV delta R8.2 and pMD2.G were gifts from Didier Trono (Addgene plasmid # 12263; http://n2t.net/addgene:12263; RRID:Addgene_12263 and Addgene plasmid # 12259; http://n2t.net/addgene:12259; RRID:Addgene_12259). Lentiviral production involved co-transfection of transfer, packaging, and envelope plasmids into human embryonic kidney (HEK 293T) cells using Lipofectamine 3000 according to the manufacturer’s instructions (ThermoFisher Scientific, MA, USA). Cell supernatant containing lentivirus was collected at 24 and 48 hours, filtered through a 0.45 μm filter, concentrated using a Lenti-X Concentrator (TakaraBio), and stored at -80°C prior to transduction.

### Cell Culture and Engineering

Human Jurkat T cells (clone E6-1) and human CD19^+^ Nalm6 lymphoblastic leukemia cells (clone G5) were purchased from ATCC (VA, USA). Cells were grown in RPMI-1640 medium (Wisent Bioproducts, QC, Canada) supplemented with 10% (v/v) Fetal Bovine Serum (FBS) and 5% (v/v) Antibiotic-Antimycotic at 37°C in 5% CO2. The absence of mycoplasma contamination in cell cultures was frequently validated using the MycoAlert Mycoplasma Detection Kit (Lonza, NY, USA).

The generation of engineered T cells involved the initial transduction of naïve Jurkat cells with the pHR_5x Gal4 UAS SEAP pPGK BFP lentivirus with 8 μg/mL polybrene for 6 hrs. Cells that expressed BFP were deemed “RE only” cells and were sorted using a FACSAria III fluorescence-activated cell sorter (BD Biosciences, CA, USA). Sequentially, RE only cells were transduced with pHR_PGK_antiCD19_synNotch_Gal4VP64 lentivirus, again with 8 μg/mL polybrene for 6 hrs. Cells that were both Myc- and BFP-positive were sorted to obtain “SynNotch+RE” cells. The expression of each engineered component was validated pre- and post-sort by staining Jurkat cells with an anti-Myc antibody (#2233S, NEB, MA, USA), followed by performing flow cytometry on a FACSCanto (BD Biosciences). All flow cytometry results were analyzed using FlowJo v10 software (FlowJo LLC, BD Biosciences).

As SEAP expression is activatable, T cells were also engineered with lentivirus to constitutively express zsGreen and Gaussia Luciferase (GLuc), the latter of which is also a secreted reporter protein detectable in culture media. This allowed the relative number of live cells in culture to be assayed at the same time as SEAP.

We used CRISPR/Cas9 to generate CD19^-^ Nalm6 cells, as previously described^28^.

### In vitro validation

CD19-targeted SynNotch+RE, RE only, and naïve Jurkat cells were co-cultured with either CD19^-^ or CD19^+^ Nalm6 cells. Within each well of a 96-well round bottom plate, 10^5^ T cells were seeded with equal number of Nalm6 cells in a total volume of 125 μL media per well. Co-cultures were then centrifuged for 5 min at 400×g to encourage cells to come into proximity. Prior to collecting media on each day, plates were spun down at 400×g for 5 min, then 120 μL of media was collected from each well, fresh media was added, and plates were centrifuged again. The collected supernatant was centrifuged at 10,000×g for 10 min and stored at -20°C until assayed. SEAP activity in supernatant (25 μL) was measured using the Great EscAPe SEAP Chemiluminescence Assay kit 2.0 (Clontech, Fremonet, CA). The GLuc activity in supernatant (20 μL) was measured using the Gaussia Luciferase Assay reagent (Targeting Systems, San Diego, USA). The luminescence signal from both assays was measured using the Glomax 20/20 luminometer from Promega (Madison, WI).

### In vivo evaluation in tumor model

All animal procedures were performed as approved by the University Council on Animal Care at the University of Western Ontario (Protocol #2020-025) and follow the Canadian Council on Animal Care (CCAC) and Ontario Ministry of Agricultural, Food and Rural Affairs (OMAFRA) guidelines. For all animal work, 4-6 weeks old female NOD.Cg-Prkdc^scid^ Il2rg^tm1WjI^/SzJ (NSG) mice were obtained from an in-house breeding colony at Western University. Tumors were initiated by subcutaneously injecting 10^6^ CD19^+^ or CD19^-^ Nalm6 cells in 50 μL of PBS and 50 μL of Matrigel (Corning, NY, USA) into the right flank of each mouse. Tumor volumes were periodically calculated using caliper measurements and the following formula:

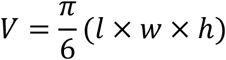

Once tumors reached ∼150 mm^3^ (∼3-5 weeks post inoculation), mice received an intratumoral injection of 10^7^ SynNotch+RE T cells suspended in 100 μL of PBS (Day 0). On days -1, 2, 4, and 7 post-delivery of immune cells, 70 μL of blood was collected from the saphenous vein from each mouse and stored in blood-collection tubes containing heparin and gel-barrier for anticoagulation and plasma separation (Becton Dickinson, ON, CA). Immediately following the blood collection, each blood sample was centrifuged at 10,000×g for 10 min to isolate blood plasma. SEAP assays were performed on 25 μL of the isolated plasma for each mouse, following the SEAP assay protocol described above.

### Histology and Immunostaining

At the study endpoint, mice were euthanized using an overdose of isoflurane. Each mouse was pressure perfused via the left ventricle using a solution of 4% paraformaldehyde (PFA). Tumors were carefully excised, submerged in 4% PFA for a period of 24 hours, and subsequently stored in PBS at 4 °C before being prepared for sectioning and staining. To facilitate sectioning, the tumors underwent a series of sucrose gradients ranging from 10% to 30% and were then frozen using optimal cutting temperature medium (Sakura Finetek). Ten-micron sections were obtained, fixed in 4% PFA for 10 minutes at room temperature, and stained with DAPI. Fluorescence images of zsGreen (T cells) and DAPI were acquired using an EVOS FL Auto 2 microscope (ThermoFisher).

### Statistics

All statistical analysis was performed using GraphPad Prism 9.0 software (GraphPad Software Inc, CA, USA). For *in vitro* cell proliferation assay and SEAP activity, a two-way ANOVA followed by Tukey’s multiple comparisons test was used. For quantifying total *in vitro* SEAP activity over 4 days, area under the curve calculations were performed. A one-way ANOVA followed by Tukey’s multiple comparisons test was used to assess GLuc activity across samples. In the case of the *in vivo* SEAP assay, a two-way ANOVA followed by Tukey’s multiple comparisons test was used for analysis. Differences between groups in the *in vivo* SEAP assay data, represented as area under the curve, were measured using an unpaired t-test. A nominal p-value of less than 0.05 was considered significant for all statistical analyses.

## RESULTS

### Engineering of immune cells with synNotch blood reporter system

Naïve T cells were first engineered with a response element (RE) encoding the secreted reporter gene SEAP (RE only cells), and then a subset of these were sequentially engineered with a synNotch receptor targeted to CD19 (SynNotch+RE cells) (Figure 1B and C). Following cell sorting, 98% of engineered cells expressed their intended components, no notable differences in BFP was observed between RE only and synNotch+RE populations (Figure 1C). No significant difference in proliferation rates over 48 hours were found when comparing naïve, RE and SynNotch+RE T cells (Figure 1D).

### SEAP activation via synNotch receptor based on antigen-dependent cell-cell interactions

To first evaluate our system, SEAP activity in media over time was measured in co-cultures of SynNotch+RE, RE only, or naive T cells with CD19^+^ or CD19^-^ Nalm6 leukemia cells at an effector:target (E:T; T cell:cancer cell) ratio of 1:1 (Figure 2). SEAP activity was significantly higher in CD19^+^ versus CD19^-^ co-cultures at both 24 and 48 hours (*p*<0.0005), and SEAP was elevated to the same degree at both time points (Figure 2C). Co-cultures of CD19^+^ or CD19^-^ Nalm6 cells with RE Only or naïve T cells did not show any significant increases in SEAP activity (Figure 2A & B). Compared to these other control conditions, a marginal but non-significant increase in SEAP activity was seen when SynNotch+RE cells were co-cultured with CD19^-^ Nalm6 cells.

**Figure 2.**
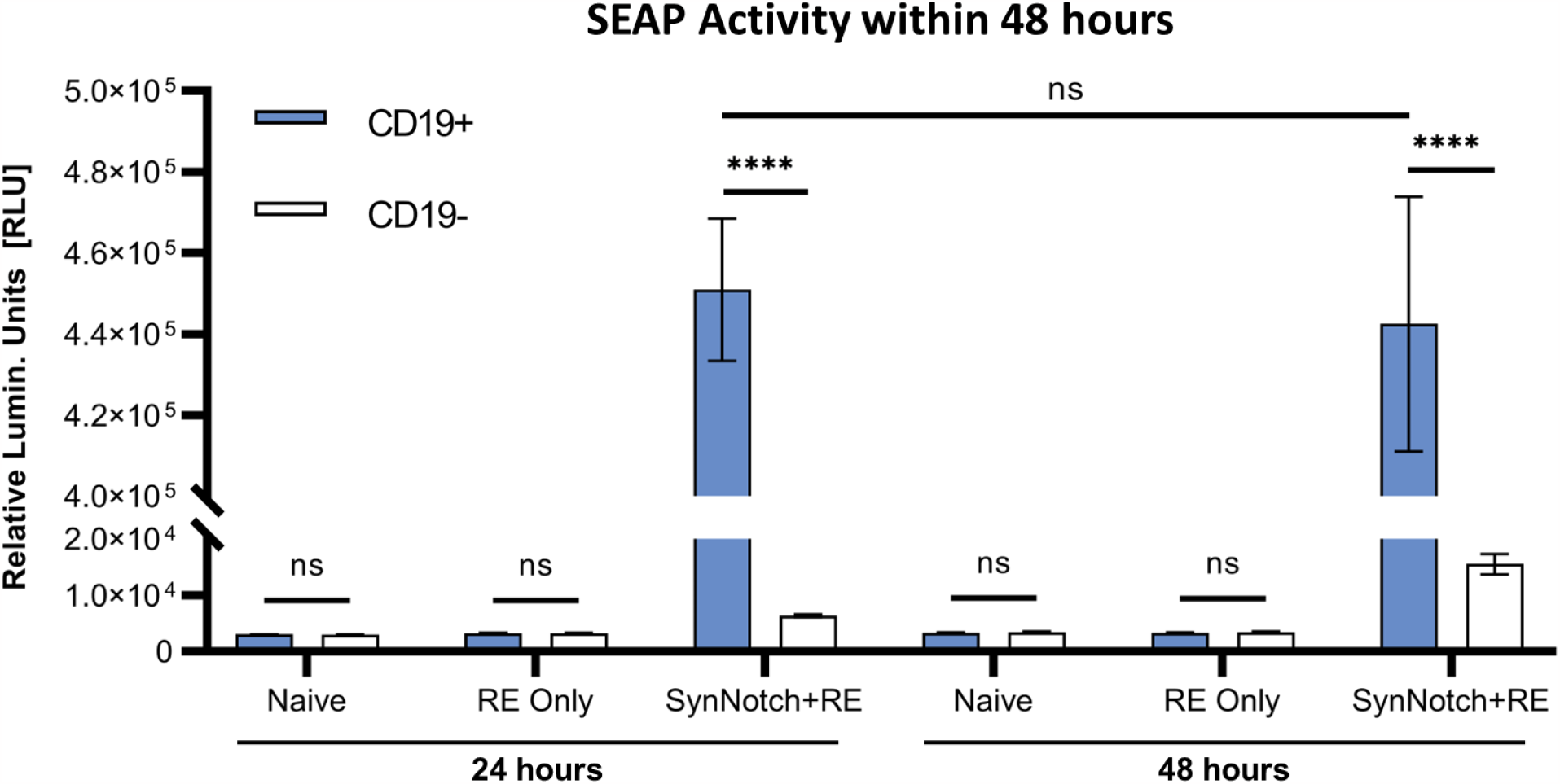
*In vitro* assessment antigen-specificity of synNotch secreted reporter system. SEAP activity values obtained from co-culturing naïve, RE only, and SynNotch+RE T cells with CD19^+^ and CD19^-^ cells at 1:1 ratio (n=3) at 24 hours and 48 hours respectively, **** represent *p*<0.0005, ns represent No Significant difference. Data are presented as mean ± SD.

Next, to model the sensitivity of this system, we varied the E:T ratio for co-cultures of SynNotch+RE cells with CD19^+^ Nalm6 cells and assessed SEAP activity over 4 days (Figure 3). First, we kept a constant number of target Nalm6 cells (10^5^ cells) and looked at the impact of increasing the number of effector cells in the co-cultures on SEAP activity in media (Figure 3A). T cells constitutively expressed GLuc, and the increases in effector cell number were confirmed via increasing GLuc activity measurements in media. Cumulative SEAP activity also increased linearly with the number of effector T cells (R^2^=1.0), with SEAP activity plateauing beyond 5×10^4^effector cells (an E:T of 0.5:1; Figure 3A). In opposition, when increasing the number of target Nalm6 cells while keeping effector numbers constant (10^5^ cells; the GLuc activity remained constant), the total SEAP activity increases with greater target number, however this time plateauing at 10^4^ target cells (an E:T of 1:1; Figure 3B).

**Figure 3.**
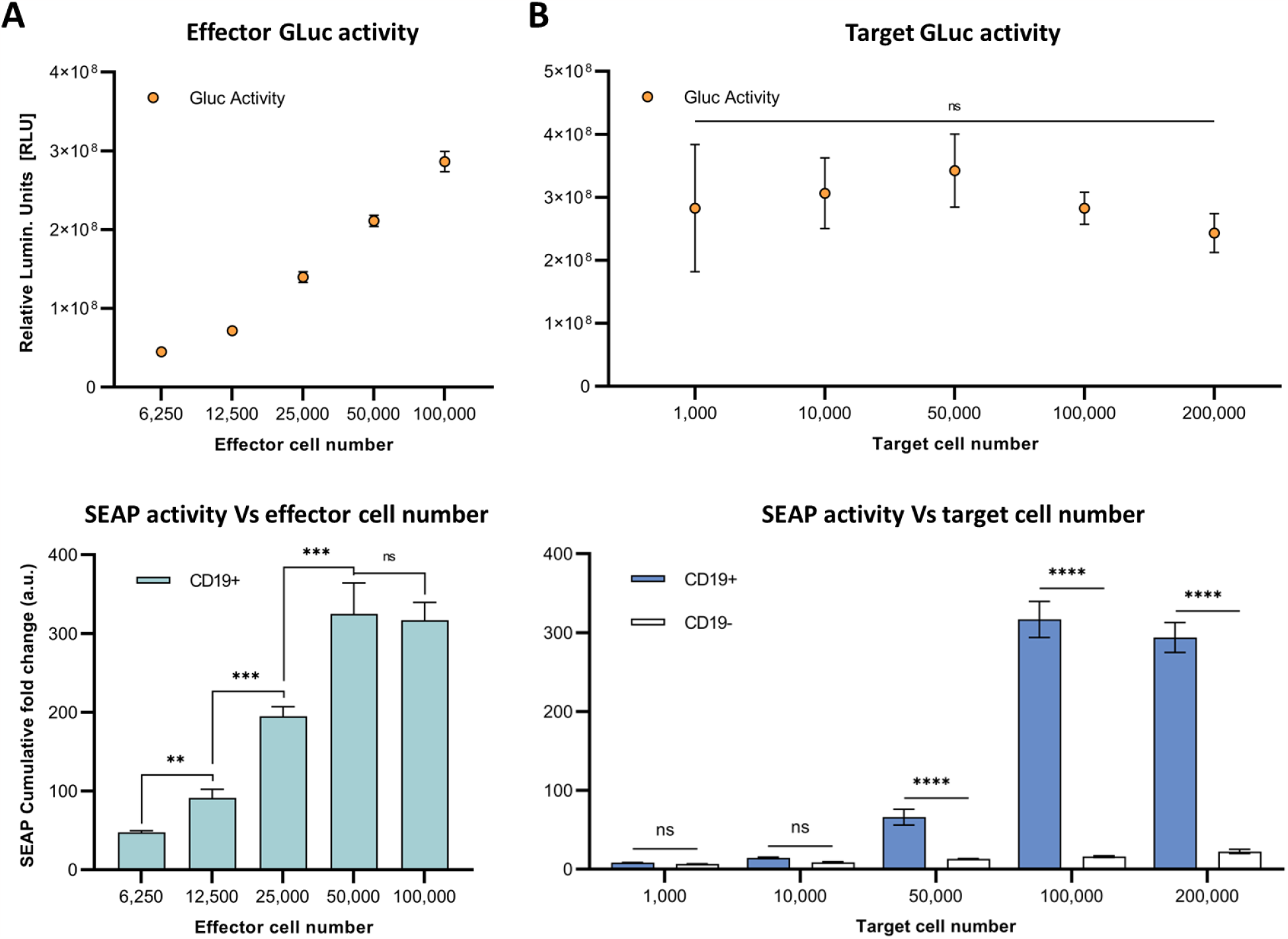
*In vitro* assessment of synNotch secreted reporter system at varying effector and target cell ratios. **(A)** Respective GLuc and SEAP activity at varying effector cell numbers. **(B)** Respective GLuc and SEAP activity at varying target cell numbers. Data presented as cumulative GLuc activity measured as the relative luminescence units (Top). SEAP activity obtained from SynNotch+RE T cells (effector cells) co-cultured with CD19^+^ cells (target cells) at varying effector cell numbers. Data presented as cumulative SEAP activity fold changes over 4 days (Bottom). Data are presented as mean ± SD (n=3). (\*\*\*\**p*<0.0005, \*\*\**p*<0.001, \*\**p*<0.01, \**p*<0.05, ns for No Significant difference).

### Antigen-dependent synNotch activation in tumors results in increased blood SEAP activity

To evaluate the synNotch system *in vivo*, subcutaneous CD19^+^ or CD19^-^ Nalm6 tumors were formed on the right flank of 4-6 weeks old female nod-scid-gamma mice. Once tumors were ∼150 mm^3^, 10^7^ SynNotch+RE T cells were injected intratumorally, blood samples were collected, and SEAP activity in blood was measured over time (Figure 4A). Prior to T cell delivery, SEAP activity was at assay background levels for both CD19^+^ and CD19^-^ groups (Figure 4B). At 48 hours post-T cell injection, mice carrying CD19^+^ tumors showed significantly elevated blood SEAP activity compared to their CD19^-^ counterparts (*p*<0.05). SEAP activity at 96 and 168 hours in the CD19^+^ cohort decreased significantly from 48 hours but remained significantly higher that from the CD19^-^ cohort (Figure 4B). TO measure the total SEAP production over 7 days, area under the curve measurements of SEAP activity over time was calculated from the longitudinal data in Figure 4B and was significantly higher for the mice bearing CD19^+^ versus CD19^-^ tumors (*p*<0.01; Figure 4C).

**Figure 4.**
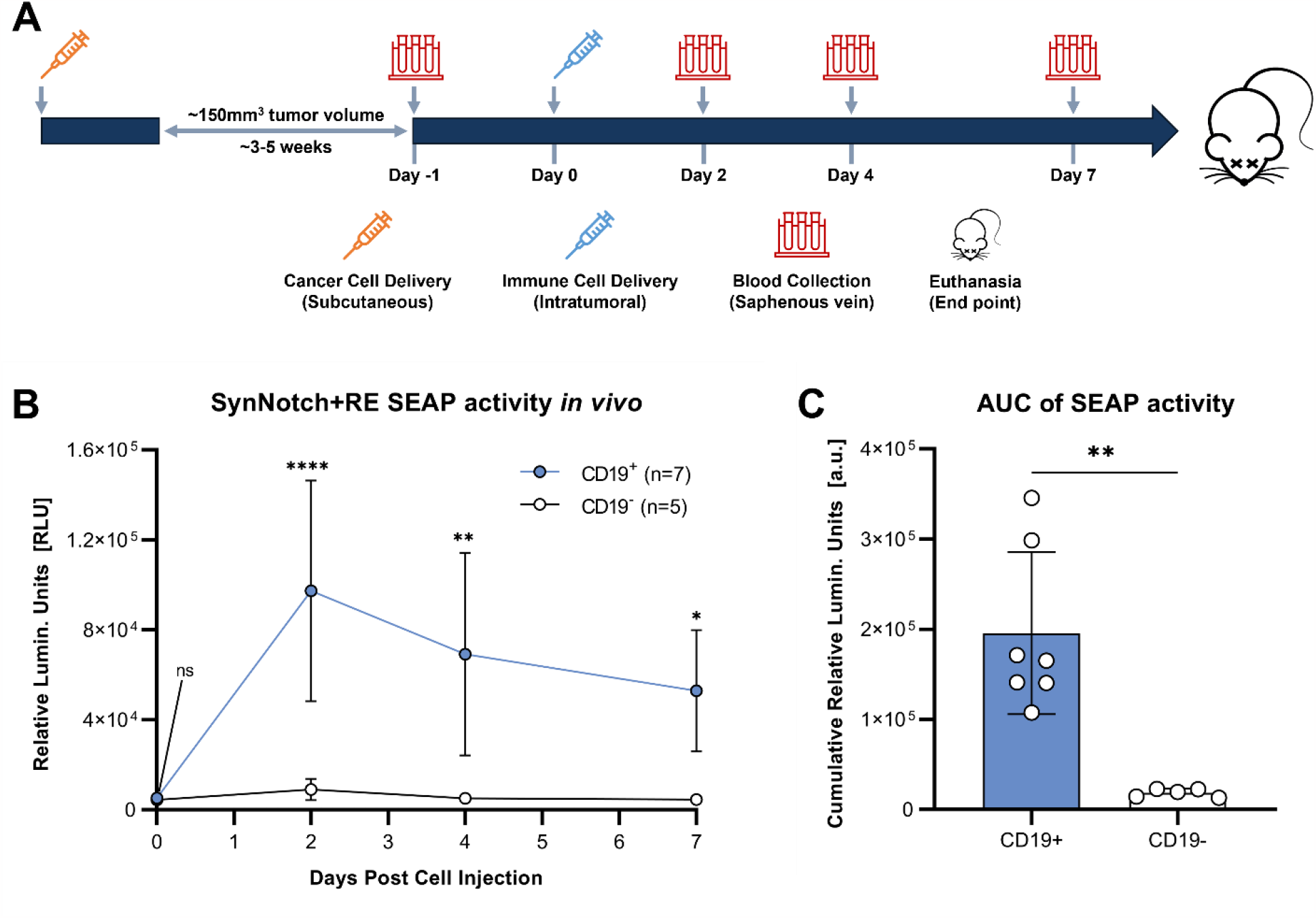
Evaluation of synNotch blood reporter system in mouse cancer model. **(A)** Experimental timeline for tumor establishment, T cell injection, and blood collection. **(B)** Average blood SEAP activity measured from mice bearing CD19^+^ (n=7) or CD19^-^ (n=5) tumors before and after the delivery of SynNotch+RE T cells. **(C)** Cumulative blood SEAP activity over 7 days (area under curve: AUC) between CD19^+^ and CD19^-^ cohorts. Data are presented as mean ± SD (\*\*\*\**p*<0.0005, \*\**p*<0.01, \**p*<0.05, ns for No Significant difference).

At endpoint, fluorescence microscopy was utilized to visualize zsGreen-positive T cells. Qualitatively, a similar number of zsGreen cells were observed in both tumor cohorts (Figure 5). This finding suggests that differences in SEAP expression were not attributable to insufficient delivery or survival of the engineered T cells across tumor types. Instead, it provides evidence supporting the persistence of the T cells post-delivery for up to 7 days.

**Figure 5.**
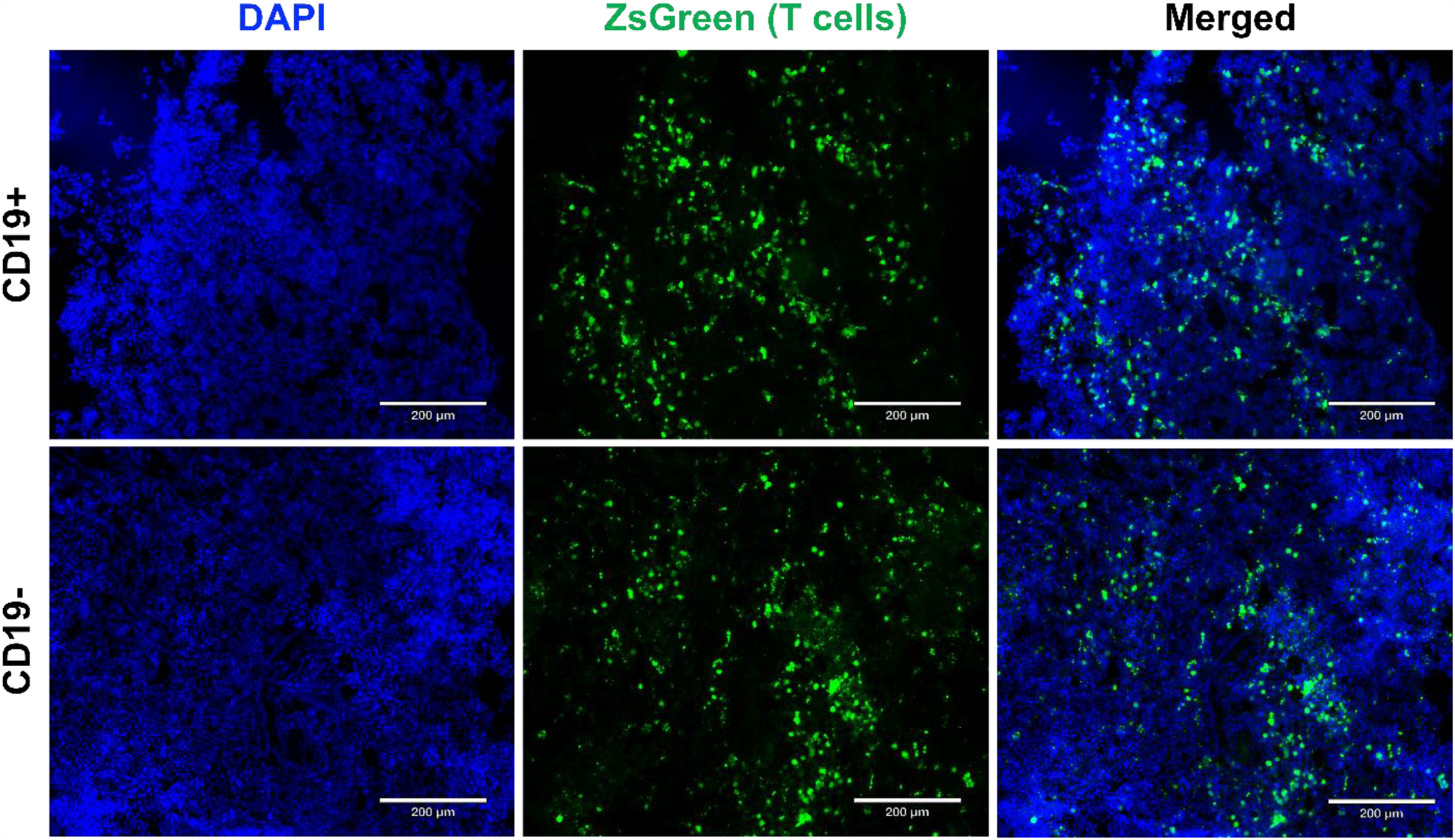
Endpoint tumor histology to visualize reporter-expressing cells. Fluorescence microscopy images (10x magnification) showcase CD19^+^ and CD19^-^ Nalm6 tumors containing SynNotch+RE T cells expressing ZsGreen. Sections were counterstained with DAPI to visualize cell nuclei. The scale bar in the images corresponds to 200 μm.

## DISCUSSION

It is attractive to utilize and customize a cell’s ability to translate extracellular triggers into the intracellular signals, as this would allow for the development of better diagnostics, therapeutics, and theranostics in patients^27,38,39^. Cell-cell communication plays a critical role in cellular immunotherapies, such as CAR-T cells, which rely heavily on interactions of the CAR with target antigens to carry out their therapeutic effects. Unfortunately, unintended interactions between immune cells and antigens on normal tissues can result in unwanted side effects and reduce treatment efficacy^40,41^. Moreover, heterogeneity in antigen density exists, where some cancer cells express the target antigens at a density that is lower than required to induce T cell cytotoxicity^42,43^. To address these issues and improve cellular therapies, it is crucial to understand adoptive cell behavior in a non-invasive manner. In this study, we developed an antigen-activatable secreted reporter system that allows for the detection of direct immune-cancer cell interactions within tumors via detectable signals in the blood.

We engineered human T cells with an activatable blood reporter system, utilizing the synNotch receptor system expressing SEAP in the response element. This system consists of two essential components: the CD19-directed receptor and the secreted reporter SEAP, with the latter being activated by the release of the transcription activator Gal4-VP64 upon antigen binding. To evaluate the engineered system, we tested its leakiness and target-dependent activatable nature. Through co-culturing naïve, RE only, and SynNotch+RE T cells with CD19^+^ or CD19^-^ cancer cells, we were able to observe baseline SEAP activity in the RE only group. Even in the presence of CD19, T cells did not express detectable SEAP without the synNotch receptor. In contrast, the SynNotch+RE group demonstrated significantly elevated SEAP activity in CD19^+^ co-cultures, while CD19^-^ co-cultures exhibited minimally elevated SEAP activity. This minimal activity may be from residual CD19 from incomplete knockout of our Nalm6 cells. These results demonstrated the dependence on both the engineered receptor and target to express SEAP, and overall shows high target specificity with little system leakiness (Figure 2).

We next aimed to investigate the dynamics of SEAP activity after antigen binding in our closed in vitro system and how varying the number of target or effector cells influenced SEAP expression. First, we maintained CD19^+^ Nalm6 and SynNotch+RE T cell co-cultures using the same number of each cell type over 2 days and observed similar elevated levels of SEAP activity at both the 24- and 48-hour mark. Due to high cell density in the 96-well plates, we observed cell death due to over confluency beyond the 48 hour time point. These results provided valuable insights into the optimal SEAP activation period for our *in vivo* experiments. Given that the activation of the synNotch system depends on physical cell-cell contact, it is important to explore the impact of varying the engineered T cell and cancer cell ratio on SEAP activity. When the number of T cells were increased at a constant cancer cell number, SEAP activity rose linearly until the number of T cells was 50% the number of cancer cells (an E:T of 0.5:1). Even when more T cells were added (an E:T of 1:1) this did not increase SEAP further, suggesting that some T cells did not interact with the cancer cells due to the possible restrictions in the proximity between them. However, having double the number of target cells (an E:T of 1:2) in our co-cultures did not increase SEAP activity further. These results raise important parameters for interpreting SEAP blood levels *in vivo* that are discussed further below and will need to be explored in future studies.

We next evaluated our synNotch blood reporter system in subcutaneous Nalm6 tumors in mice. Due to the influence of E:T ratio on the amount of SEAP produced *in vitro*, we chose to inject a standard number of T cells (10^7^) into tumors of a standard volume (∼150 mm^3^). Significantly elevated blood SEAP activity was only observed in mice carrying the CD19^+^ tumor following intratumoral delivery of SynNotch+RE T cells. SEAP activity peaked at 48 hours and marginally declined thereafter up to 7 days. As the blood half-life of SEAP is 3 hours^30^, the continued elevation of SEAP over the 7 days is likely due to continued GAL4-VP64 binding to the response element after synNotch cleavage and/or additional synNotch interactions with CD19 over time. The length of RE activation after synNotch-antigen binding is unknown and could provide a better understanding of RE transgene expression over time. Fluorescence microscopy for T cells was used to confirm that the persisting SEAP signal was due to continued T cell presence in the tumor. Qualitative assessment of CD19^+^ and CD19^-^ tumors showed similar numbers of immune cells, indicating that the difference in SEAP expression between the two groups was due to the presence of the target CD19, not variations in T cell survival.

Measurements of SEAP activity were ended at 7 days due to reaching heavy tumor burden (endpoint >1.5cm^3^), however literature has reported persisting synNotch-driven CAR and luciferase expression in mice for up to 11 days following intravenous injection of T cells^44^. Compared to our findings, this could be attributed to differences in T cell persistence within the tumor between intratumoral and intravenous delivery methods^45^. We hypothesis that intratumoral delivery results in immediate activation of the synNotch receptor in the target-expressing tumor, while systemic delivery involves T cell traffic to the tumor, resulting in gradual and sustained reporter expression^46^. To fully understand the dynamics of synNotch expression and optimize its *in vivo* potential, it will be valuable to investigate the rate of replenishment of new synNotch receptors on the cell surface. Additionally, in future studies, it will be valuable to understand whether intratumoral delivery causes proteolytic cleavage of synNotch receptors at a rate that outpaces their replenishment.

SEAP is a useful reporter for monitoring immune cell activity non-invasively and can potentially provide a single measure of all the immune-cancer interactions in the whole body. However, in the context of multi-organ metastatic diseases, it is likely also important to obtain more spatial intertumoral information on the activity of immune cells. Non-invasive imaging tools like MRI and PET-based reporter genes can be used for this purpose as we and another group have recently demonstrated^28,29^. Future work to build response elements that encode both imaging and secreted reporter genes would allow both spatial information to be acquired with infrequent and relatively expensive imaging, and more frequent whole-body monitoring with relatively low-cost blood tests.

A significant limitation of the detection system described in this study is the challenge of accurately reporting the live T cell and cancer cell numbers *in vivo*. While the system demonstrated its ability to specifically report the amount of local immune-cancer cell communications, translating the use of GLuc as a measurement of live T cells *in vivo* was a major obstacle. Several reasons contribute to this limitation, particularly the reliability of detecting GLuc within blood samples due to its short half-life, and the clearance rate of GLuc through urine, making it difficult to establish consistent output^47^. Additionally, it would be valuable to monitor cancer cell numbers simultaneously with T cell numbers. Although the Jurkat T cells used in this study lacked cytotoxic abilities to focus on establishing a detection system, monitoring cancer cell numbers would be crucial in future studies where this system is combined with other cell-based therapies. Estimating cancer cell numbers through tumor volume measurement may not be accurate in future studies involving immunocompetent mice due to the infiltration of various immune cells^48^.

An alternative and more optimal preclinical approach would be to incorporate different imaging reporter genes into both T cells and cancer cells during the engineering process. Imaging reporters like bioluminescence imaging (BLI), MRI-contrast, and PET reporter genes have been previously applied in the study of many cell-or gene-based therapies^49–51^. Among these options, BLI is frequently applied in pre-clinical small animal studies of cell-based therapies^51^. By using different imaging reporters, we can overcome the limitations associated with using GLuc and improve our ability to accurately monitor and quantify live T cells and cancer cells *in vivo*. This advancement will significantly enhance the utility and effectiveness of the detection system for future studies and potential therapeutic applications.

The activatable secreted reporter system described in the study offers a novel approach to measure local immune-cancer cell interactions beyond conventional methods *in vivo*. It can potentially aid in the development and monitoring of novel cell therapies by allowing non-invasive and rapid interrogation of cell behavior. Moreover, when coupled with other imaging systems, it has the potential to provide insights into the reasons for the failure of certain cell-based immunotherapies in patients. Additionally, this system can be broadly applicable in preclinical research, enabling the study of cellular behavior during development, normal physiology, and disease progression. Overall, it represents a promising tool that can advance cell-based therapies and contribute to more effective and targeted treatments in the future.

## ACKNOWLEDGEMENTS

This research received funding provided by the Canadian Institutes of Health Research, led by J.A.R. (Grant #202104PJT-462838). We extend our gratitude to Dr. John Kelly and Dr. Ying Xia for their invaluable contributions to animal procedures and the generation of CD19-Nalm6 cells. Special thanks also go to Dr. Kristin Chadwick, the manager of the London Regional Flow Cytometry Facility, for her expert guidance in cell sorting.

## AUTHOR CONTRIBUTIONS

Y.F., T.W., and J.A.R. designed research; Y.F., and T.W. performed research; Y.F. contributed new reagents/analytic tools; Y.F., T.W., and J.A.R. analyzed data; J.A.R. provided supervision; and Y.F., T.W., and J.A.R. wrote the paper.

